# Universal and quantitative detection of double-stranded RNAs as a signature of pan-virus infections using a luciferase-based biosensor

**DOI:** 10.1101/2025.04.07.647538

**Authors:** Michihito Sasaki, Eri Fujii, Satoko Sasaki, Takuma Ariizumi, Kei Konishi, Akihiko Sato, William W. Hall, Hirofumi Sawa, Yasuko Orba

## Abstract

Viral infections produce double-stranded RNA (dsRNA) during replication, which trigger host innate immune responses. Immunoassays using anti-dsRNA antibodies have been widely employed to detect viral dsRNA. In this study, we used a luciferase-based dsRNA biosensor which consists of protein kinase R (PKR)-derived dsRNA binding domains fused to split luciferase subunits. Here, we demonstrate the use of the dsRNA biosensor to measure viral dsRNA in RNA specimens extracted from cells infected with Japanese encephalitis virus (JEV). Moreover, the biosensor reacts to a broad-spectrum of dsRNAs from infection with representatives of various viral families including positive- and negative-sense single-stranded RNA (ssRNA) viruses, dsRNA viruses, and DNA viruses.

We validated the specific interaction between the dsRNA biosensor and viral RNA including subgenomic flavivirus RNA (sfRNA) through RNA immunoprecipitation. Additionally, we observed luminescence signals directly from lysates of JEV-infected cells after cell lysis and phase separation with Triton X-114. Finally, we used the biosensor to assess the activity of antiviral compounds. In summary, our results demonstrate that the luciferase-based dsRNA biosensor offers a simple, homogeneous, and high-throughput platform for quantifying viral replication, presenting a promising alternative to antibody-based dsRNA detection methods.

## Introduction

The genomes of viruses include encapsided DNA, single-stranded RNA (ssRNA), or double-stranded RNA (dsRNA) and viruses are classified based on their genome types and replication strategies (Baltimore classification)(1). dsRNA comprises the nucleic acid of dsRNA viruses; however, this type of nucleic acid is also produced by ssRNA viruses as a replication intermediate and as a byproduct during viral replication in cells(2). As positive-sense ssRNA viruses replicate, they generate both viral genome−antigenome hybrid dsRNA (replicative form) and partial dsRNA that has multiple transcribing genomic RNAs (replicative intermediate) (3, 4). Copy-back defective viral genomic RNA is a partial viral genomic RNA characterized by reverse complementary 5′ and 3′ ends forming a dsRNA stem structure and is generated during the replication of various non-segmented negative-sense ssRNA viruses(5). In the case of DNA virus infection, converging bidirectional transcription of the viral genome produces complementary mRNA transcripts that can form dsRNA(6, 7). Collectively, the aberrant accumulation of dsRNA serves as a biomarker for viral infection.

Host cells counter viral infection through several dsRNA sensor proteins such as RIG-I-like receptors (RLRs), protein kinase R (PKR), and oligoadenylate synthases (OASes)(2). In particular, PKR is a dsRNA-dependent Ser/Thr protein kinase consisting of two tandem dsRNA binding domains (dsRBD) and a kinase domain. PKR recognizes dsRNA in a sequence-independent manner and requires a minimum of 30 to 33 bp of dsRNA for its activation(8, 9). When PKR interacts with dsRNA, the former is autophosphorylated, followed by the phosphorylation of eukaryotic initiation factor 2 alpha (eIF2α) which results in translational shutoff and viral proliferation(10, 11). PKR can sense various viral infections, and it mediates a universal antiviral effect in host cells. In response, some viruses have evolved strategies to antagonize the innate immune responses mediated by the dsRNA-PKR-eIF2α axis(10, 11).

Anti-dsRNA monoclonal antibodies specifically recognize dsRNA longer than 40 bp in a sequence-independent manner(12). Clone J2, the most commonly employed anti-dsRNA monoclonal antibody, has been used as a tool to detect a wide range of viral infections in various immunoassays including immunofluorescence(3, 13, 14), immunohistochemistry(15, 16), immunoprecipitation(17), and immunochromatographic assays(18). J2 has also been used to determine the subcellular localization and accumulation of dsRNA during the process of viral genome replication; furthermore, it has been used to study the mechanisms of antiviral innate immune responses in various viral infections(3, 14). Virological studies have predominantly relied on immunoassays using anti-dsRNA antibodies to detect and quantify dsRNA; however, these assays have limitations in sensitivity, quantitation and throughput. Therefore, it is desirable to develop other more sophisticated dsRNA detection methods.

One alternative dsRNA detection tool is currently available as a luciferase assay system using a dsRNA biosensor (Lumit dsRNA Detection Assay, Promega). This system is based on a split luciferase complementation using small and large subunits of NanoLuc luciferase, referred to as smBiT and LgBiT, respectively, by the manufacturer(19). The biosensor consisting of PKR-derived dsRBD fused with smBiT or LgBiT (dsRBD-smBiT or dsRBD-LgBiT, respectively) binds to dsRNA and reconstitutes the dsRBD-tagged smBiT and LgBiT subunits as a functional NanoLuc luciferase(20, 21). The performance of this luciferase assay system has been evaluated by quantifying artificially synthesized dsRNA and detecting dsRNA contamination as a byproduct in *in vitro* transcription products(20). The affinity of PKR for virus-induced dsRNA facilitates the use of this luciferase-based dsRNA biosensor as a new research tool in the field of virology. In this study, we have characterized the interaction between this biosensor and viral RNA and examined its utility in various types of viral infections.

## Results

### Detection of dsRNA accumulation produced by flavivirus infection using the luciferase-based dsRNA biosensor

We firstly used Japanese encephalitis virus (JEV), a single-stranded positive-sense RNA virus in the family *Flaviviridae*, to examine whether the dsRNA biosensor can detect viral replication in cells. Previous studies have used immunostaining with anti-dsRNA monoclonal antibodies to show the accumulation of dsRNA in cells infected with flaviviruses(22, 23). Consistent with previous reports, our immunofluorescence analysis with anti-dsRNA antibody J2 demonstrated the accumulation of dsRNA in BHK-21 cells infected with JEV (Fig. 1A). Total RNA was extracted from JEV-infected cells at different time points, and quantitative RT-PCR (qRT-PCR) analysis revealed increases in intracellular viral RNA levels over time (Fig. 1B). We also observed a time-dependent increase of dsRNA in Quantitative ELISA with J2 antibody, while no clear dsRNA signal was detected at 24 hpi (Fig. 1C). To examine the reactivity of dsRBD-LgBiT and dsRBD-smBiT with dsRNA produced during viral replication, RNA samples from JEV-infected cells were incubated with dsRBD-LgBiT and dsRBD-smBiT, and the resulting luminescence was measured (Fig. 1D). The results showed higher luminescence signals in the RNA samples at 24 (*p*=0.053), 48 (*p*<0.0001) and 72 hpi (*p*<0.0001) than that in RNA of uninfected controls (Fig. 1E). Next, we performed the luciferase assay and qRT-PCR using RNA samples mixed with RNAs from JEV-infected and uninfected cells in different ratios. The luminescence and qRT-PCR signals were closely correlated (*R^2^* = 0.9829, Fig. 1F), although we observed background luminescence in the assay using RNA from uninfected cells, which affected the limit of detection of the assay. The luminescence signals from JEV-infected Vero and N2a cells indicated that the assay is independent of cell type (Fig. 1G and 1H). Overall, these results indicate the applicability of the luciferase-based dsRNA biosensor for detecting dsRNA accumulation during JEV replication.

**Figure 1.**
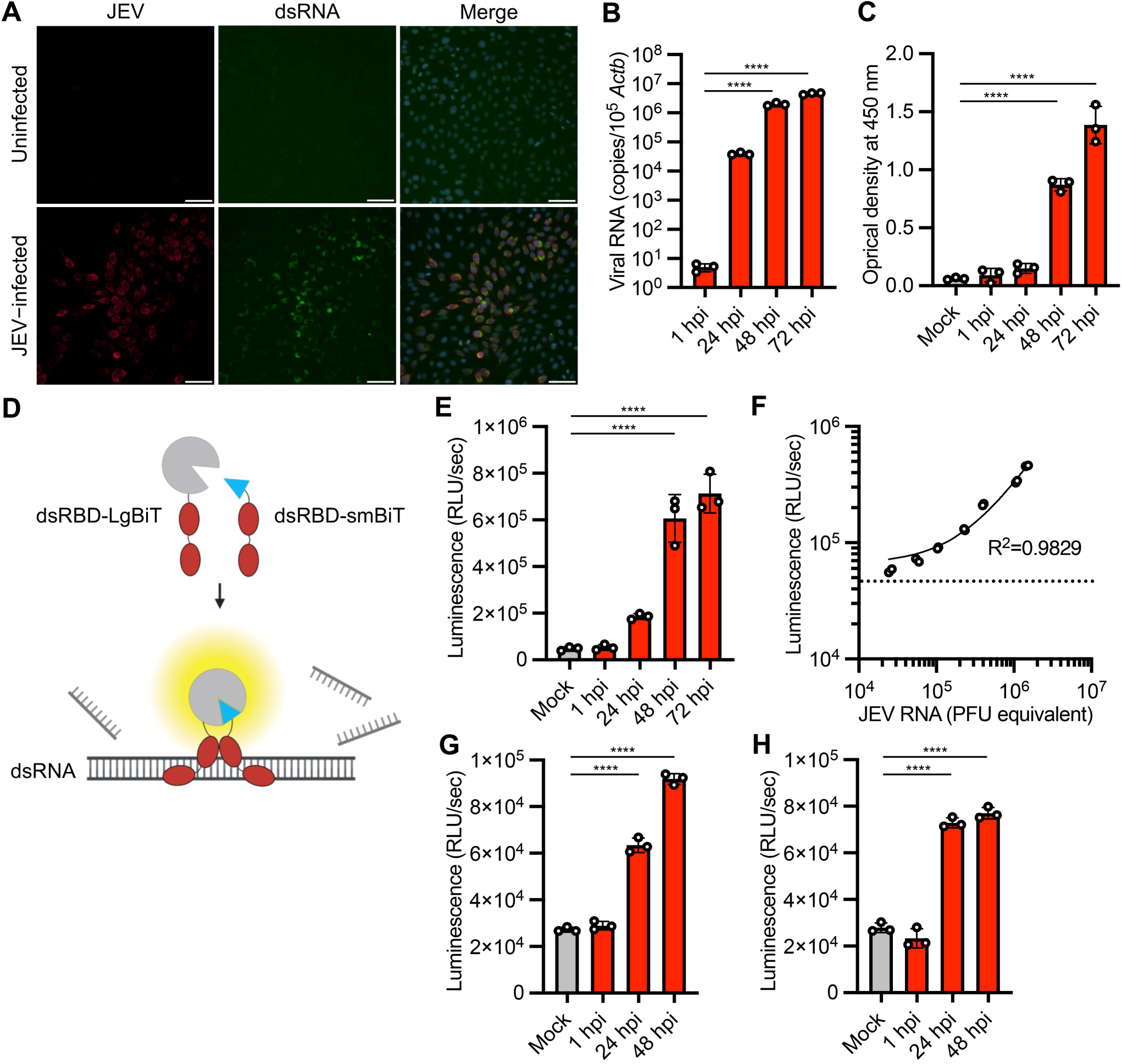
Quantitative detection of dsRNA in JEV infection by luciferase assays with the dsRNA biosensor. **(A)** Immunofluorescence staining of BHK-21 cells infected with JEV or mock-infected cells at 24 hpi. Cells were stained with anti-JEV (red), anti-dsRNA (green), and Hoechst 33342 (blue). **(B)** Viral RNA loads in cells at the indicated time points of JEV infection were analyzed by qRT-PCR. qRT-PCR data were normalized to endogenous *Actb*. **(C)** dsRNA loads in cells at the indicated time points of JEV infection were analyzed by J2 antibody-based ELISA. **(D)** Schematic representation of dsRNA detection in RNA sample by split nanoluciferase complementation assay using the dsRNA biosensor (dsRBD-smBiT and dsRBD-LgBiT). **(E)** Total RNA was extracted from BHK-21cells infected with JEV and analyzed by luciferase assay with the dsRNA biosensor. **(F)** Correlation of the qRT-PCR targeting JEV RNA and luciferase assays with the dsRNA biosensor. RNA samples from JEV-infected and uninfected BHK-21 cells were mixed at different ratios. The RNA mixtures were analyzed by qRT-PCR and luciferase assay to measure the levels of JEV RNA and dsRNA, respectively. The horizontal dashed line indicates the luminescence value of uninfected cells in the assay (the detection limit of dsRNA related to JEV infection). *R^2^* indicates the linear correlation coefficients between qRT-PCR and dsRNA detection assays. **(G, H)** Total RNA was extracted from Vero (G) and N2a (H) cells infected with JEV and analyzed by luciferase assays with the dsRNA biosensor. The values shown are mean ± standard deviation (SD) of triplicate (B, C, E, G, H) or duplicate (F) samples. \*\*\*\**p* < 0.0001 by one-way ANOVA with Dunnett’s test.

### Broad-spectrum reactivity of the luciferase-based dsRNA biosensor to different viral infections

To examine whether the dsRNA biosensor can detect the replication of various virus types, we tested this in infections caused by the following viruses: positive-sense ssRNA chikungunya virus (CHIKV, family *Togaviridae*) and Severe acute respiratory coronavirus 2 (SARS-CoV-2, family *Coronaviridae*); negative-sense ssRNA rabies virus (RABV, family *Rhabdoviridae*) and La Crosse virus (LACV, family *Peribunyaviridae*); dsRNA rotavirus A (RVA, family *Sedoreoviridae*) and Nelson Bay orthoreovirus (NBV, family *Spinareoviridae*); and dsDNA mpox virus (MPXV, family *Poxviridae*) and herpes simplex virus 1 (HSV-1, family *Orthoherpesviridae*). Immunofluorescence analysis with anti-dsRNA and anti-viral protein antibodies demonstrated dsRNA accumulation in viral antigen-positive Vero cells infected with CHIKV, LACV, NBV and MPXV, Vero-TMPRSS2 cells infected with SARS-CoV-2, BHK-21 cells infected with RABV, and MA104-T2T11D cells infected with RVA (Fig. 2A). dsRNA luciferase assay of total RNA extracted from cells showed 9.5-fold to 22.3-fold increases in luminescence signal in RNA samples from cells infected with CHIKV, SARS-CoV-2, RABV, LACV, RVA, NBV and MPXV compared with uninfected cells (Fig. 2B). In contrast, we failed to detect immunofluorescence signal of dsRNA and observed limited (2.0-fold) increase in luminescence signal in Vero cells infected with HSV-1 (Fig. 2A and 2B), consistent with previous reports that virion host shutoff protein of HSV-1 destabilizes dsRNA and limits its accumulation in cells(6, 7). These results suggest that the dsRNA biosensor detects the infection with a broad range of virus families through the accumulation of dsRNA in the infected cells.

**Figure 2.**
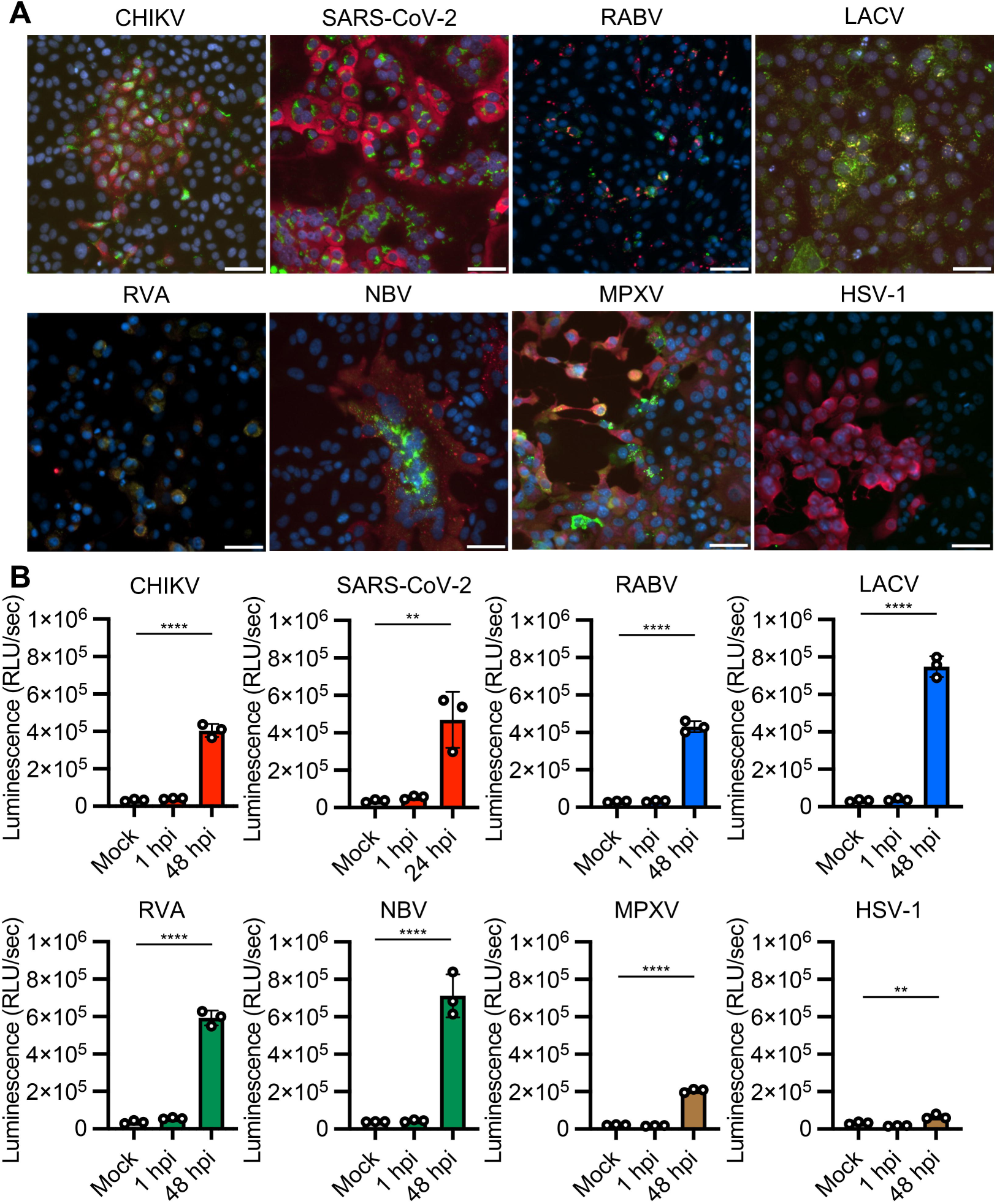
Application of the dsRNA biosensor to different viral infections. Positive-sense ssRNA viruses [Chikungunya virus (CHIKV), SARS-CoV-2], negative-sense ssRNA viruses [Rabies virus (RABV), La Crosse virus (LACV)], dsRNA viruses [Rotavirus A (RVA), Nelson Bay orthoreovirus (NBV)], DNA viruses [Mpox virus (MPXV), and Herpes simplex virus 1 (HSV-1)] were inoculated onto monolayers of Vero (CHIKV, LACV, NBV, MPXV and HSV-1), Vero-TMPRSS2 (SARS-CoV-2), BHK-21 (RABV), and MA104-T2T11D cells (RVA). **(A)** Immunofluorescence staining of the infected cells with Hoechst 33342 nuclear dye (blue), antibodies targeting dsRNA (green) and virus proteins (red). Scale bars, 50 μm. **(B)** Luciferase assay with the dsRNA biosensor was used to measure dsRNA in RNA extracts from mock-infected and virus-infected cells. The values shown are mean ± SD of triplicate samples. \*\**p* < 0.01, \*\*\*\**p* < 0.0001 by one-way ANOVA with Dunnett’s test.

### Specificity of the interaction of luciferase-based dsRNA biosensor with flavivirus genomic and subgenomic RNAs

We investigated the target of the dsRNA biosensor in RNA samples from JEV infected cells. First, we compared the reactivity of the dsRNA biosensor to RNA samples with or without heat denaturation, which resolves double-stranded structure of dsRNA and secondary and tertiary structures of ssRNA. Heat denaturation of sample RNA reduced the luminescence signal of JEV-infected cells to a baseline level, similar to that of uninfected cells (Fig. 3A), indicating that the viral infection-specific increase of the luminescence signal depends on the RNA structure. We next examined the nature of the interaction between the dsRNA biosensor and JEV RNA by RNA immunoprecipitation (RIP) with RNA from JEV-infected cells using dsRBD-LgBiT, anti-LgBiT antibody, and protein G beads. The LgBiT protein lacking the dsRBD domain was used as a control in the RIP assay. Immunoblotting results showed the successful capture of dsRBD-LgBiT and LgBiT on the Protein G beads (Fig. 3B). Viral genome RNA, viral mRNA, and subgenomic flavivirus RNA (sfRNA) derived from the 3′-UTR are synthesized during active viral replication in cells(24). Quantification of the amount of viral RNA in the immunoprecipitate by qRT-PCR targeting the NS2A region and 3′-UTR of the JEV genome revealed the specific interaction between the viral RNA and dsRBD-LgBiT (Fig. 3C and 3D). To determine the type of viral RNA in the immunoprecipitate, we performed northern blot analysis with an RNA probe targeting the 3′-UTR of the JEV genome. The denaturing gel electrophoresis followed by northern blot analysis identified the JEV sfRNA and genomic RNA including replicative intermediate and replicative form in the RNA samples extracted from JEV virions and JEV-infected cells (Fig. 3E). Furthermore, the analysis of the immunoprecipitants revealed that dsRBD-LgBiT binds to not only the JEV RNA genome but also sfRNA (Fig. 3E). To validate the specific interaction between dsRBD-LgBiT and sfRNA, we performed additional RIP assay with *in vitro* transcribed sfRNA (525 nt in length). Sequence scrambled sfRNA (525 nt in length), 525 and 3000 nt ssRNAs of partial coding sequence (CDS) regions of JEV genome [CDS(525) and CDS(3000)], and 717 nt ssRNA of AcGFP complete CDS were also synthesized by *in vitro* transcription and used as ssRNA controls for the RIP assay. The synthesized sfRNA, but not control ssRNAs, was enriched in the immunoprecipitate with anti-LgBiT and dsRBD-LgBiT (Fig. 3F). Collectively, these results suggest that the dsRNA biosensor specifically recognizes viral RNA including sfRNA synthesized in the cells infected with JEV.

**Figure 3.**
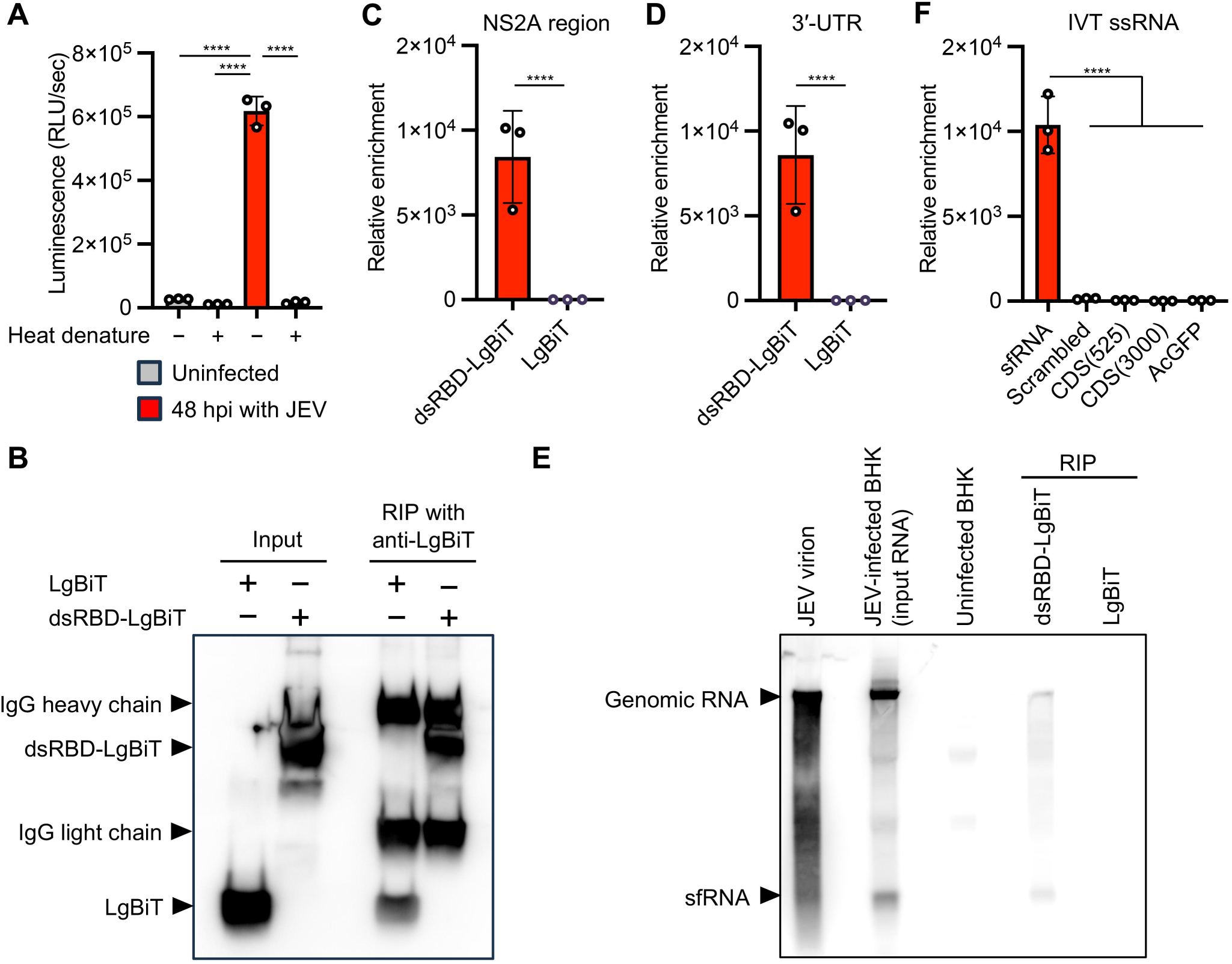
Target specificity of the dsRNA biosensor. **(A)** Effect of heat denaturation on dsRNA sensing by the dsRNA biosensor. Heat-denatured and non-denatured RNA samples were analyzed by luciferase assays with the dsRNA biosensor. **(B-D)** RNA immunoprecipitation (RIP) assays with total RNA extracted from BHK-21 cells at 48 hpi with JEV using dsRNA biosensor dsRBD-LgBiT and anti-LgBiT antibody. LgBiT protein without dsRBD was used as a control in the RIP assay. Immunoprecipitates were analyzed by immunoblotting with anti-LgBiT (B) and qRT-PCR targeting the JEV NS2A gene (C) and the 3′-UTR of the JEV genome (D). **(E)** Northern blotting of the immunoprecipitates using digoxigenin DIG–labeled RNA probe targeting the 3′-UTR of the JEV genomic RNA including replicative form and replicative intermediate. RNA extracted from the culture supernatant containing JEV virions, JEV-infected and uninfected BHK-21 cells were included in the blotting to estimate the size of the JEV genome and subgenomic flavivirus RNA (sfRNA). **(F)** RIP assays with *v* ssRNAs of JEV sfRNA, sequence-scrambled sfRNA (scrambled), partial coding sequences of JEV genome [CDS(525) and CDS(3000)], and complete coding sequence of AcGFP. Relative enrichment of each ssRNA was calculated by dividing the ssRNA amounts of immunoprecipitates using dsRBD-LgBiT with those of immunoprecipitates using LgBiT protein control. The values shown are mean ± SD of triplicate samples. \*\*\*\**p* < 0.0001 by one-way ANOVA with Tukey’s test (A, F) or two-tailed Welch’s *t*-test (C, D).

### Quantitative detection of dsRNA in a homogeneous cell lysates using the luciferase-based dsRNA biosensor

All the above experiments for dsRNA detection were performed with RNA extracted from cells. We aimed to simplify the protocol by skipping the RNA extraction process and detecting viral dsRNA directly in cell lysates using the dsRNA luciferase assay. First, we assessed whether detergent contaminating the cell lysate could affect the dsRNA luciferase assay. Using an extracted RNA sample containing 1% of conventional detergents, we found that contamination by all tested detergents decreased the luminescence signal from the dsRNA biosensor (Fig. 4A). Next, we suspended JEV-infected cells in lysis buffer with different concentrations of digitonin and added the dsRNA biosensor to the mixture. The luminescence signal from the dsRNA biosensor was accentuated by adding high concentrations of digitonin in the cell lysis buffer, but the signal peaked at 0.25% of digitonin and decreased at 1% and 4% of digitonin, probably due to the carry-over of digitonin from the input cell lysate containing high concentrations of the detergent to the dsRNA luciferase assay mixture (Fig. 4B). To reduce detergent carry-over after sufficient cell lysis, we employed a phase separation approach using Triton X-114. Triton X-114 is a nonionic detergent that forms a homogeneous solution on ice, but the solution separates into aqueous and detergent-enriched phases above 20 (25). We suspended JEV-infected cells in the lysis buffer with Triton X-114 on ice and then incubated the cell lysate at 37 , followed by brief centrifugation to separate the phases (Fig. 4C). The dsRNA luciferase assay detected a high luminescence signal in the aqueous phase of the cell lysate even with high concentrations of Triton X-114. Notably, the phase separation procedure significantly increased the luminescence signals from cell lysates containing 0.25%, 1% and 4% of Triton X-114 (Fig. 4D). Overall, these results demonstrate that phase separation of cell lysates with Triton X-114 enable us to obtain the RNA fraction with low detergent concentrations and is a simple and reproducible way to directly detect dsRNA in cells.

**Figure 4.**
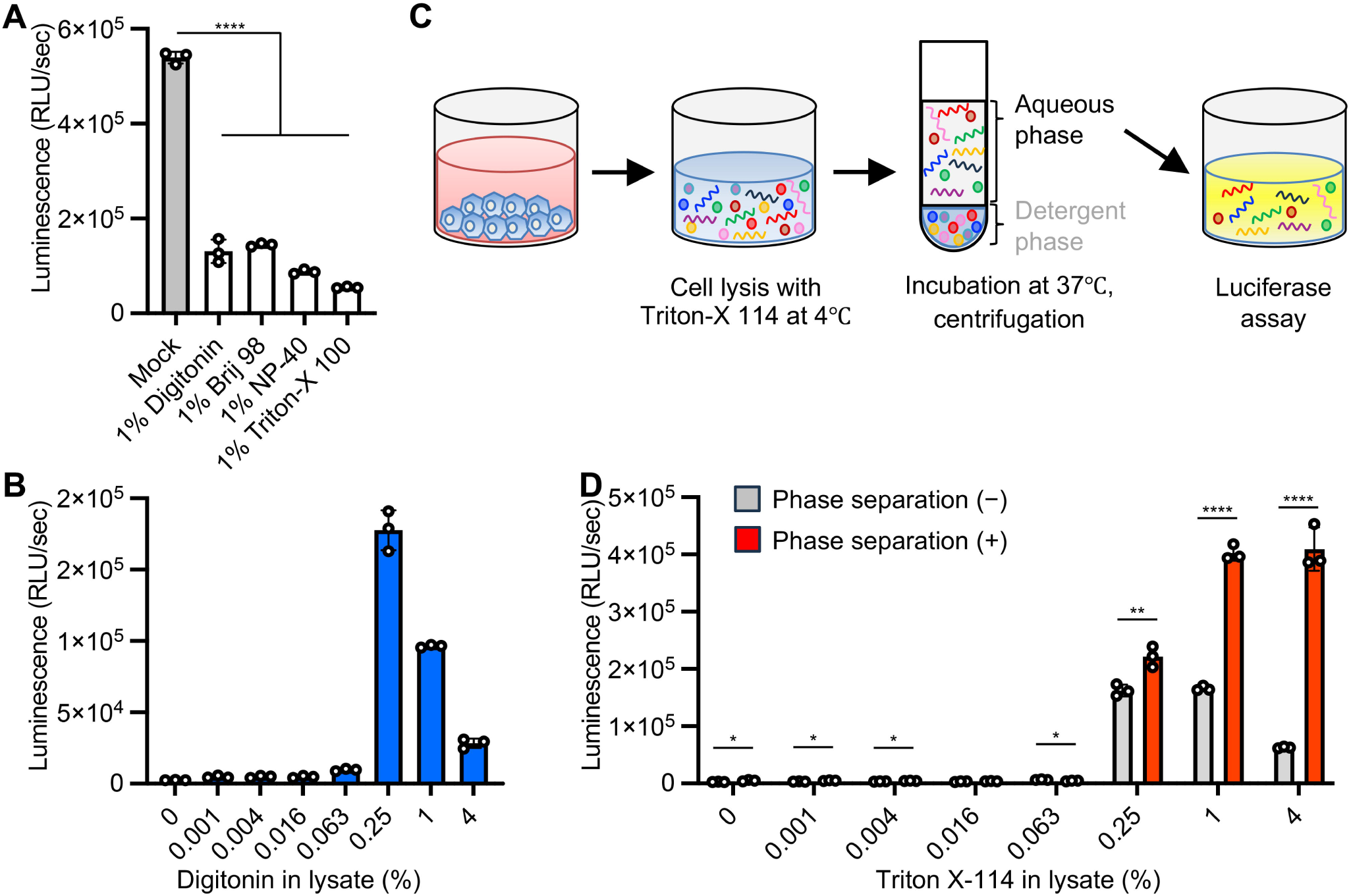
Detection of dsRNA in cell lysates using the dsRNA biosensor. **(A)** RNA samples extracted from JEV BHK-21 cells at 48 hpi with JEV were mixed with the indicated detergents at a final concentration of 1% and then analyzed by the luciferase assay for dsRNA detection with the dsRNA biosensor. **(B)** Cells at 48 hpi following JEV infection were suspended in lysis buffer containing different concentrations of digitonin and analyzed by the luciferase assay with the dsRNA biosensor. **(C)** Schematic representations of cell lysis and phase separation with Triton X-114. The aqueous phase of the cell lysate was used for the luciferase assay with the dsRNA biosensor. **(D)** Cells at 48 hpi with JEV were suspended in lysis buffer containing different concentrations of Triton X-114. Cell suspensions containing Triton-X114 [phase separation (-)] and the aqueous phase of the cell lysate after centrifugation [phase separation (+)] were analyzed by the luciferase assay with the dsRNA biosensor. The values shown are mean ± SD of triplicate samples. \**p* < 0.05, \*\**p* < 0.01, \*\*\*\**p* < 0.0001 by one-way ANOVA with Dunnett’s test (A) or two-tailed Student’s *t*-test (D).

We next examined whether our assay can measure the activity of antiviral compounds. Ribavirin and NITD008 are nucleotide analogs that act as antiviral agents against flaviviruses, including JEV(26–28). Both immunofluorescence staining and qRT-PCR assays demonstrated the anti-JEV activities of ribavirin and NITD008 (Fig. 5A-5D). Consistent with the results of conventional assays, the cell-directed dsRNA luciferase assays showed that JEV infection decreased in a dose-dependent manner in response to these antivirals (Fig. 5E and 5F). These results suggest that the cell-directed dsRNA luciferase assay is a simple and fast method to estimate the magnitude of viral replication and the activity of antiviral compounds.

**Figure 5.**
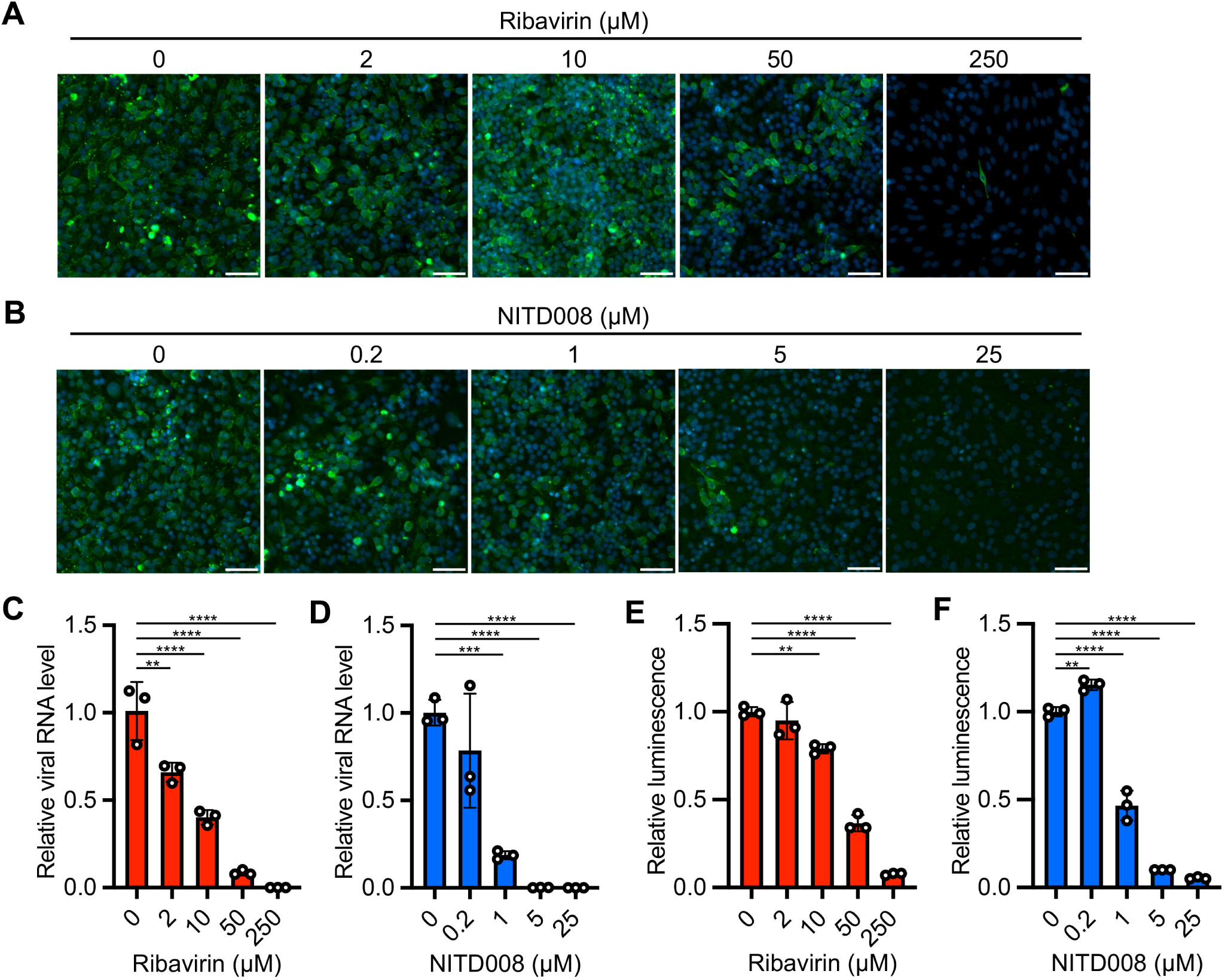
Application of the dsRNA biosensor to antiviral assays (A, **B)** Cells were infected with JEV in the presence of different concentrations of ribavirin (A) or NITD008 (B). Immunofluorescence staining of the cells at 48 hpi using anti-JEV (green) and Hoechst 33342 (blue). Scale bars, 50 μm. **(C, D)** Viral RNA levels were measured in the cells at 48 hpi by qRT-PCR. Relative viral RNA amounts in the presence of ribavirin (C) and NITD008 (D) were compared with those in the absence of the antivirals. Data were normalized to endogenous *Actb*. **(E, F)** Cells at 48 hpi with JEV were lysed in Triton X-114-containing lysis buffer. The aqueous phase of the lysates was obtained after phase separation and used for dsRNA quantification with the luciferase assays with the dsRNA biosensor. Relative luminescence signals in the presence of ribavirin (E) and NITD008 (F) were compared with those in the absence of the antivirals. The values shown are mean ± SD of triplicate samples. \*\**p* < 0.01, \*\*\**p* < 0.001, \*\*\*\**p* < 0.0001 by one-way ANOVA with Dunnett’s test (C-F).

## Discussion

Replication of RNA and DNA viruses in cells produces viral genome-derived RNA duplexes that stimulate the host antiviral immune responses through dsRNA sensors such as RLRs, TLR-3, and PKR. In this study, we used the dsRNA biosensor dsRBD-LgBiT and dsRBD-smBiT that employ the NanoLuc luciferase complementation process to detect viral dsRNA in infected cells. We could demonstrate that luminescence signal from the biosensor increases in a time-dependent manner and correlates with the viral RNA load in JEV-infected cells. The biosensor could also be used to detect different viral infections including those caused by ssRNA viruses, dsRNA viruses, and DNA viruses, without the need for antibodies and oligonucleotide primers targeting specific viral proteins and genomes. These results indicate that the luciferase-based dsRNA biosensor is capable of estimating the magnitude of various types of viral infections including emerging and as yet undiscovered viruses.

Anti-dsRNA monoclonal antibodies have been widely used to detect viral dsRNA in various immunoassays as the protocol has broad applications and involves a simple workflow. However, antibody-based immunoassays can sometimes be associated with nonspecific reactions and variable signal intensity(29). The dsRNA luciferase assay is not an immunoassay; rather, the dsRNA biosensor carries the dsRBD from PKR. Both the dsRNA antibody and PKR recognize dsRNA in a sequence-independent manner. Using a crosslinking and immunoprecipitation sequencing (CLIP-seq) analysis, Kim *et al*. revealed that PKR as well as the anti-dsRNA antibody J2 can interact with various types of dsRNA(30). Compared with the conventional immunoassays, the dsRNA luciferase assay with the dsRNA biosensor offers a simple, rapid and high-throughput workflow to detect dsRNA in RNA samples. However, this assay has been developed for dsRNA detection in the extracted RNA samples and is susceptible to the presence of high amount of ssRNA and detergent in the specimens. The results from our studies suggest that the dsRNA luciferase assay is a versatile method for viral dsRNA detection with quantification and would be applicable to sequence-independent detection of known and unknown viral infections in cells.

PKR displays sequence-nonspecific interaction and autoactivates by not only dsRNA but also ssRNA with an imperfect short stem flanked by single-stranded tails(31, 32). The dsRNA biosensor recognizes dsRNA larger than 30 bp regardless of the nucleotide sequence composition(20). Although we were able to demonstrate the interaction between the biosensor and the JEV genome and sfRNA, the details on the target regions within the viral RNAs remain elusive. There are multiple stem-loop structures in the 5′- and 3′-UTR of the flavivirus genome and sfRNA(33, 34). Notably, a previous study showed that viral sfRNA induces the activation of host PKR in cells(35). In addition, the long dsRNA structure of flavivirus RNA genome consisting of a hybrid of the 5′-UTR and 3′-UTR was formed during genomic RNA cyclization (a process that occurs during flavivirus replication)(33, 34). We thus speculate that the stem-loops in JEV genomic RNA and sfRNA are putative binding targets of dsRNA biosensor.

In this study, we showed the applicability of the dsRNA biosensor in a homogeneous cell lysate. Although the interaction between the dsRNA biosensor and dsRNA was attenuated by common detergents used for cell lysis, implementing a phase separation step following cell lysis with Triton X-114 can effectively mitigate detergent interference while maintaining the assay’s sensitivity. Our cell-directed dsRNA luciferase assay was applicable to measure the antiviral activity of compounds from cell lysates, and the results correlated well with those of qRT-PCR. Implementing the dsRNA luciferase assay directly on cells is a simple and fast way to evaluate the magnitude of viral infections. We believe that the assay will also be useful for high-throughput screening of antiviral compounds and host factors.

Although our study demonstrated the broad reactivity of the dsRNA biosensor in the detection of various viral infections, it has the following limitations. First, the target dsRNAs in each viral infection, except for JEV, remain elusive. The dsRBD in the biosensor is derived from PKR; thus, identifying the targets of the biosensor will contribute to our knowledge on the process of PKR activation in response to viral infection. Second, the presence of endogenous dsRNA needs to be considered. The cellular levels of host-derived dsRNAs such as inverted Alu repeats, bidirectional transcription of mitochondrial DNA, and cellular mRNA with an extended 3′-UTR increase in response to pathophysiological conditions(30, 36). In addition to virus-induced dsRNA, host-derived dsRNAs are also sensed by endogenous dsRNA sensors. In neurons, mRNA with an extended 3′-UTR acts as an endogenous dsRNA that triggers constitutive type I IFN production, conferring an intrinsic antiviral state(37). However, we assumed that the endogenous dsRNA had little effect in this study based on the result of heat denaturation treatment of total RNA from uninfected cells, which showed limited impact on the luciferase-based dsRNA assay. Finally, we observed background luminescence even in total RNA samples from uninfected cells. This indicates that it may be necessary to further optimize the luciferase assay protocol to ensure its wide dynamic range and robustness in future applications.

## Materials and methods

### Cell lines

BHK-21 (RCB1423, RIKEN BRC) and C6/36 cells (CRL-1660, ATCC) were maintained in Eagle’s Minimum Essential Medium (MEM) containing 10% fetal bovine serum (FBS). N2a cells (IFO50081, JCRB) were maintained in MEM containing 10% FBS and nonessential amino acids (Gibco). Vero-TMPRSS2 cells stably expressing human TMPRSS2 were established in our laboratory as described previously(38, 39).

Vero-TMPRSS2, Vero (CRL-1586, ATCC), and 293T cells (632180, Takara Bio) were maintained in Dulbecco’s Modified Eagle’s Medium (DMEM) containing 10% FBS. MA104-T2T11D cells expressing human TMPRSS2 and TMPRSS11D were established in our laboratory as described previously(40) and maintained in MEM containing 10% FBS and 10% tryptose phosphate broth (BD Difco).

### Viruses

JEV (Mie/41/2002 strain)(41), CHIKV (SL10571 strain)(42), SARS-CoV-2 (WK-521 strain)(43), RABV (HEP strain)(44), NBV (MB/07 strain)(45) and MPXV (Zr-599 strain)(46) were provided by the National Institute of Infectious Diseases, Japan. HSV-1 (F strain) was provided by the Institute of Medical Science, University of Tokyo, Japan. RVA (SA11 strain) and LACV (VR-1834) were obtained from the ATCC. Working virus stocks were prepared by passaging through C6/36 cells for JEV; BHK-21 cells for RABV; Vero cells for CHIKV, NBV, MPXV, LACV and HSV-1; Vero-TMPRSS2 cells for SARS-CoV-2; and MA104-T2T11D cells for RVA. The working virus stocks were titrated by standard plaque assay (JEV, CHIKV, NBV, MPXV, LACV, HSV-1 and SARS-CoV-2) or focus assay (RABV and RVA)(40, 47).

### Immunofluorescence staining

BHK-21 cells on 24 well plate (Corning) were inoculated with 100 μl of inoculum containing JEV or RABV at a multiplicity of infection (MOI) of 0.1 for 1h and then cultured in MEM containing 2% FBS for 24 h. Vero cells on 24 well plate were respectively inoculated with 100 μl of inoculum containing CHIKV, LACV, NBV, or HSV-1 at an MOI of 0.1 and then cultured in DMEM containing 2% FBS for 24 h. Other subset of Vero cells on 24 well plate were inoculated with 100 μl of inoculum containing MPXV at an MOI of 0.01 for 1h and then cultured in DMEM containing 2% FBS for 48 h. Vero-TMPRSS2 cells on 24 well plate were inoculated with 100 μl of inoculum containing SARS-CoV-2 at an MOI of 0.1 for 1h and then cultured in DMEM containing 2% FBS for 24 h. MA104-T2T11D cells on 24 well plate were inoculated with 100 μl of inoculum containing RVA at an MOI of 0.1 for 1h and then cultured in MEM containing 2% FBS and 10% tryptose phosphate broth for 24 h. Cells were fixed at 24 h post-infection (hpi) except for MPXV, which was fixed at 48 hpi with 3.7% buffered formaldehyde, followed by permeabilization with 0.5% Triton X-100 in PBS for 5 min and double–staining with anti-dsRNA mouse IgG2a monoclonal antibody (clone J2; 10010200, SCICONS) and virus-specific primary antibody in 25% Block Ace (KAC) in PBS for 1 h. We used anti-CHIKV E1 mouse IgG1 monoclonal (MAB12424, Abnova), anti-SARS-CoV-2 nucleocapsid rabbit monoclonal (GTX635679, GeneTex), anti-RABV nucleoprotein mouse IgG2b monoclonal (3R7-4G4, HyTest), anti-LACV mouse IgG2b monoclonal (MA1-10801, Thermo Fisher Scientific), anti-RVA goat polyclonal (AB1129, Merck), anti-Vaccinia virus rabbit polyclonal (cross reactive with MPXV; PA17258, Thermo Fisher Scientific), anti-HSV-1 rabbit polyclonal (B0114, DAKO), anti-JEV rabbit polyclonal (serum from JEV-immunized rabbit in our laboratory), and anti-NBV rabbit polyclonal (serum from NBV-immunized guinea pig in our laboratory) antibodies(48) for the viral antigen staining. The cells were subsequently stained for 30 min with the following appropriate secondary antibodies: anti-mouse IgG2a Alexa Fluor 488-conjugated goat polyclonal (A21131, Thermo Fisher Scientific), anti-mouse IgG1 Alexa Fluor 594-conjugated goat polyclonal (A21125, Thermo Fisher Scientific), anti-mouse IgG2b Alexa Fluor 594-conjugated goat polyclonal (A21145, Thermo Fisher Scientific), anti-rabbit IgG Alexa Fluor 594-conjugated goat polyclonal (A11037, Thermo Fisher Scientific), anti-guinea pig IgG Alexa Fluor 647-conjugated goat polyclonal (A21450, Thermo Fisher Scientific), anti-mouse IgG Alexa Fluor 488-conjugated donkey polyclonal (A21202, Thermo Fisher Scientific), and anti-goat IgG Alexa Fluor 568-conjugated donkey polyclonal (A11057, Thermo Fisher Scientific). Cell nuclei were stained with Hoechst 33342 (Thermo Fisher Scientific). Fluorescent images were captured using an IX73 fluorescence microscope (Olympus).

### RNA measurement by qRT-PCR

Virus growth kinetics were determined with BHK-21 cells infected with JEV at an MOI of 0.01. Total RNA was extracted from the cells at different time points with the Direct-zol RNA MiniPrep Kit (Zymo Research) and analyzed by qRT-PCR with the EXPRESS One-Step Superscript qRT-PCR Kit (Thermo Fisher Scientific) and a primer/probe set targeting JEV *NS2A*(49). The hamster *Actb* gene was used as an internal control for normalization(50). For the RIP assay, *in vitro* transcribed (IVT) ssRNAs and the 3′-UTR of the JEV genome in the immunoprecipitates were quantified, by qRT-PCR on the extracted RNA using a One-Step TB Green PrimeScript PLUS RT-PCR Kit (Takara Bio). JEV RNA levels were calculated by the standard curve, comparative ΔCt or ΔΔCt methods. The probe and primers used in this study are summarized in Table S1 (Supplementary materials).

### Detection of dsRNA in RNA extracts

BHK-21 cells on 24 well plate were infected with JEV or RABV at an MOI of 0.01.

Vero cells on 24 well plate were individually infected with CHIKV, LACV, NBV, MPXV, and HSV-1 at an MOI of 0.01. Vero-TMPRSS2 cells and MA104-T2T11D cells on 24 well plate were infected with SARS-CoV-2 and RVA, respectively, at an MOI of 0.01. At 1 and 48 hpi, the cells were lysed with 300 μl of RNAiso Plus (Takara Bio). Total RNA was extracted from the cell lysates with Direct-zol RNA MiniPrep Kit , with mock-infected cells included as controls. The extracted RNA samples were subjected to dsRNA luciferase assay using the Lumit dsRNA biosensor (dsRBD-LgBiT and dsRBD-smBiT) with dsRNA assay buffer and luminescence detection substrate B provided in the Lumit dsRNA Detection Assay kit (W2041, Promega) according to the manufacturer’s instructions with slight modifications. Briefly, a total of 200 to 500 ng RNA in 5 μl water was mixed and incubated with dsRBD-LgBiT and dsRBD-smBiT in 95 μl of dsRNA assay buffer for 1 h at room temperature. The mixture of RNA and biosensor was then added to 25 μl of luminescence detection substrate B diluted in dsRNA assay buffer on 96-well half area black microplates (3694, Corning). After 15-min of incubation, the luminescence was measured with a GloMax Discover Microplate Reader (Promega). In parallel with the dsRNA luciferase assay, dsRNA amount in RNA samples was also measured by ELISA with dsRNA ELISA kit, J2-based (Exalpha Biologicals) following the manufacture’s instruction. 500 ng of RNA samples were loaded in each well and the values of optical density at 450 nm were measured with iMark microplate reader (Bio-Rad).

Heat denaturation treatment of dsRNA was conducted by heating RNA samples for 5 min at 65 , then chilling on ice for 3 min, heating for 2 min at 95 , and finally chilling on ice for 3 min. The heat denatured RNA samples was subjected to the dsRNA luciferase assay as described above.

### dsRNA detection in the cell lysates

BHK-21 cells on a 24-well plate were infected with JEV at an MOI of 0.01. At 48 hpi, cells were lysed in 100 μl/well of cell lysis buffer (25 mM Tris-HCl [pH 7.5], 150 mM NaCl, 5 mM EDTA) supplemented with different concentrations of digitonin (0.001 to 4%), Triton X-114 (0.001 to 4%), NP-40 (1%), Brij 98 (1%) (all from Sigma), and Triton X-100 (1%) (Nacalai Tesque) for 5 min on ice. The phases in the Triton X-114-containing cell lysates were separated by first incubating the mixture for 1 min at 37 and then centrifuging at 9000 × *g* for 30 s. The amount of dsRNA was measured in the upper detergent-free phase. We analyzed cell lysates that had both undergone and not undergone phase separation by mixing 5 μl of the lysates with dsRBD-LgBiT and dsRBD-smBiT in 95 μl of dsRNA assay buffer and analyzing the solution using the Lumit dsRNA Detection Assay kit as described above.

### RIP assay with cellular RNA samples

Fifty microliters of protein G-coupled magnetic beads (Dynabeads protein G; Veritas) were conjugated with 10 μg of anti-LgBiT monoclonal antibody (N7100, Promega) for 15 min and then incubated with 22.8 pmol of control LgBiT (Promega) or dsRBD-LgBiT in PBS with 0.002% Tween 20 (PBST) for 30 min. Then 10 μg of total RNA extracted from BHK-21 cells at 48 hpi with JEV was incubated with the magnetic beads in 500 μl of dsRNA assay buffer supplemented with 60 units of a recombinant RNase inhibitor (Takara Bio) overnight at 4 with rotation. The mixture was then washed 3 times with PBST, and 10% of the beads were suspended in SDS sample buffer [4% SDS, 100 mM Tris-HCl (pH 6.8), 20% glycerol, 10% 2-mercaptoethanol] for immunoblotting. The remaining 90% of the beads were suspended in RNAiso plus and the suspension was subjected to RNA extraction with Direct-zol-96 MagBead RNA Kit (Zymo Research). The RNA extract was analyzed by qRT-PCR for JEV RNA levels in the immunoprecipitate as described above and by Northern blotting for JEV RNA size in the immunoprecipitate.

Our immunoblotting protocol began by resolving input and immunoprecipitated proteins by SDS-PAGE, followed by blotting of the proteins onto an Immobilon-PSQ PVDF membrane (Merck). The membrane was incubated with anti-LgBiT monoclonal antibody (N7100) overnight at 4 and then incubated with HRP-conjugated anti-mouse IgG polyclonal antibody (AMI3404, BioSource) for 30 min at room temperature. Immune complexes were detected using Immobilon Western Chemiluminescent Substrate (Merck) on an Amersham ImageQuant 800 (Cytiva).

Northern blotting was performed using the DIG Northern Starter Kit (Roche) according to the manufacturer’s instructions with slight modifications. Briefly, we resolved the following on 1.2% SeaKem GTG agarose-2.2 M formaldehyde gels : RNA samples eluted from the magnetic beads, RNA extracted from BHK-21 infected with JEV, control RNAs from uninfected BHK, and JEV virions in culture supernatants. The RNAs were transferred onto a Nytran SuPerCharge nylon membrane using a Turboblotter (Cytiva) and UV crosslinked at 120 mJ/cm^2^ in an HL-2000 HybriLinker (UVP). For the digoxigenin (DIG)–labeled RNA probes, template cDNA of the 3′-UTR of the JEV genome fused with T7 promoter sequence was amplified by RT-PCR and applied to *in vitro* transcription with T7 RNA polymerase. The UV-crosslinked membrane was hybridized with DIG–labeled RNA probe in ULTRAhyb Ultrasensitive Hybridization Buffer (Thermo Scientific).

Hybridization complexes were detected with alkaline phosphatase-conjugated anti-DIG antibody and CDP-Star (all from Roche) on Amersham ImageQuant 800.

### RIP assay with IVT ssRNA samples

For *in vitro* transcription, DNA templates at nucleotide positions 10441 to 10965 for of sfRNA, positions 9871 to 10395 for CDS (525) and positions 7396 to 10395 for CDS (3000) in JEV genome (GenBank accession no. AB241119) were prepared and fused with T7 promoter sequence by RT-PCR with PrimeScript II High Fidelity One Step RT-PCR Kit (Takara Bio). DNA template of sequence-scrambled sfRNA with T7 promoter was synthesized through gBlocks gene fragment synthesis (IDT). DNA template of AcGFP CDS with T7 promoter was prepared by PCR with PrimeSTAR Max DNA polymerase and pAcGFP1-C1 vector (Takara Bio). Sequence information of scrambled sfRNA and primers for PCR/RT-PCR was summarized in Table S2 and S3 (Supplementary materials). IVT ssRNAs were prepared by *in vitro* transcription with Takara IVTpro mRNA Synthesis Kit (low dsRNA) (Takara Bio) and then purified with NucleoSpin RNA Clean-up XS kit (Macherey Nagel). For RIP assay, 3.5 μl of protein G-coupled magnetic beads were conjugated with 0.5 μg of anti-LgBiT monoclonal antibody (N7100) for 15 min and then incubated with 1.4 pmol of control LgBiT or dsRBD-LgBiT in PBST for 30 min. The magnetic beads were incubated with 10 ng of IVT ssRNA samples in 300 μl of dsRNA assay buffer supplemented with 10 units of a recombinant RNase inhibitor and 3000 ng of total RNA from uninfected BHK-21 cells for 2h at 4 with rotation. The mixtures were then washed 3 times with PBST and suspended in RNAiso plus. The RNA was extracted with Direct-zol-96 MagBead RNA Kit and analyzed by qRT-PCR for each ssRNA levels in the immunoprecipitates as described above.

### Evaluation of antiviral efficacy

BHK-21 cells on 24-well plates were infected with JEV at an MOI of 0.01 in the presence of different concentrations of ribavirin (Wako) or NITD008 (MedChemExpress). At 48 hpi, a subset of cells were fixed with 3.7% buffered formaldehyde and stained with anti-JEV rabbit polyclonal antibody. Another subset of cells on different 24-well plates were lysed in 100 μl/well of cell lysis buffer supplemented with 1% Triton X-114 for 5 min on ice. After phase separation, 5 μl of the upper detergent-free phase was analyzed using the Lumit dsRNA Detection Assay kit to quantify dsRNA as described above. The remaining lysate was subjected to RNA extraction and qRT-PCR as described above.

### Statistical analysis and reproducibility

Statistical significance was determined by one-way analysis of variance (ANOVA) with Dunnett’s test (Figs. 1B, 1C, 1E, 1G, 1H, 2B, 4A and 5C-5F), one-way ANOVA with Tukey’s test (Fig. 3A and 3F), two-tailed Welch’s *t*-test (Fig. 3C and 3D) and two-tailed Student’s *t*-test (Fig. 4D). All statistical tests were carried out using Prism version 9.5.1 (GraphPad software). All experiments were repeated independently 2 or 3 times with similar results. Most results are presented as mean± SD of triplicate samples as specified in the figure legends.

### Data availability

All data associated with this work are included in this article.

## Acknowledgments

We thank Dr. Saijo and Dr. Ebihara of the National Institute of Infectious Diseases, Japan for providing JEV, CHIKV, SARS-CoV-2, RABV, NBV and MPXV, Dr. Kawaguchi of Tokyo University for providing HSV-1, and Dr. Ito of Gifu University for support in the establishment of MA104-T2T11D cells. We also thank Dr. Inagawa of Promega Japan for technical support in the Lumit dsRNA detection assays.

