## Supplementary Tables for "Universal and quantitative detection of double-stranded RNAs as a signature of pan-virus infections using a luciferase-based biosensor"

### Supplementary materials

#### Table S1. Primers used for virus RNA quantification in this study

| Primer Name | Sequence (5' to 3') | Reference |
| --- | --- | --- |
| JEV NS2A-F | AGCTGGGCCTTCTGGT | (49] |
| JEV NS2A-R | CCCAAGCATCAGCACAAG | |
| JEV NS2A-Probe | FAM- CTTCGCAAGAGGTGGACGGCCA-BHQ1 | |
| Mie41(10123)-F | CAAGTCTGGAACAGGGTATGG | This study |
| Mie41(10356)-R | TCTTCTGAGGGAAGTCATGTAATC | |
| Mie41(10582)-F | CGCACCGGAAGTTGAAAGAC | |
| Mie41(10770)-R | TGTTGTTTCCACGGGGTCTC | |
| Scrambled sfRNA-F | AAAGCCACCTGTATCTAGGGTA | |
| Scrambled sfRNA-R | AGCACCGTCTCATGTTTATGG | |
| Ham_Actb-F | ACTGCCGCATCCTCTTCCT | (50) |
| Ham_Actb-R | TCGTTGCCAATGGTGATGAC | |
| Ham_Actb-Probe | FAM-CCTGGAGAAGAGCTATGAGCTGCCTGATG-BHQ1 | |

#### Table S2. Primers used for ITV template DNA preparation

| Primer Name | Sequence (5' to 3') | Product |
| --- | --- | --- |
| T7-sfRNA-F | ATTTAATACGACTCACTATAGGGTGGAGTCAGGCCAG CAAAAGCTG | JEV sfRNA |
| T7-sfRNA-R | AGATCCTGTGTTCTTCCTCACCACCAGCTA | |
| T7-Scr-F | ATTTAATACGACTCACTATAGGGACCGACTCG | Scrambled sfRNA |
| T7-Scr-R | ATCTTAGTGCGGGCTTACGCTCCTCTGA | |
| T7-CDS(525)-F | ATTTAATACGACTCACTATAGGGCCGTGCAGAGGGCA GGATGAGCTGA | CDS(525) |
| T7-CDS-R | CTAAATAACCCTGTCCTCCTGAATTAATACATC | |
| T7-CDS(3000)-F | ATTTAATACGACTCACTATAGGGCTGGAAAGAACTAC TCCTTTGATGCAGA | CDS(3000) |
| T7-CDS-R | CTAAATAACCCTGTCCTCCTGAATTAATACATC | |
| T7-AcGFP-F | ATTTAATACGACTCACTATAGGGATGGTGAGCAAGGG CGCCGAGCTGT | AcGFP |
| T7-AcGFP-R | CTTGTACAGCTCATCCATGCCGTG | |

**Table. S3. Nucleotide sequence of IVT template of scrambled sfRNA**

| Name | Sequence (5' to 3') |
|---|---|
| T7-Scrambled sfRNA | ATTTAATACGACTCACTATAGGGACCGACTCGCGATCCACGTAGAAGGGCATTA AAGGCCGCGGCACCTGTATGTAGATCGTACAGACAGAAGAATTGACATAGCCA TGGTATTAAGTTGGATGGGTACGCACTCCCGGGCCCGCGGGAGGGCACCAAGA GCTACAATAGACGCAACCGGCTCTGTAAAAGCCACCTGTATCTAGGGTACCTAG GGAAGACGCACGGGCCCCACTTCGAGTACCTCTCACGCCGGCAAGGGTTTTGA TCGATGGGAAACCCGGAGTAGACGCATAGGAGGCCAGCGCGAGCACGAAGGC AAAGGCTGAGTCCCCAGCTAACCAAGGCAGCCATGTAACTTCGGCTGCCATAA ACATGAGACGGTGCTTCGCAGTCGCTTACGGGGCCCAGAACTTTCAGGCTGAA AAAACTATGTGAAGGGAGAGCAGCGTGTTTTCGGTGGATGAGGACGAGCAAG CTTGATCGGGTCCGGATCCTCGTGCCATTAGAGGAGGCCTAATCAGAGGAGCGT AAGCCCGCACTAAGAT |
